# *ProLung*™-*budesonide* Inhibits SARS-CoV-2 Replication and Reduces Lung Inflammation

**DOI:** 10.1101/2021.05.05.442779

**Authors:** Kameswari S. Konduri, Ram Pattisapu, Jogi Pattisapu, Girija G. Konduri, John Zwetchkenbaum, Bidhan Roy, Monalisa Barman, Adria Frazier, Brett L. Hurst, Nejat Düzgüneş

**Author notes:** Corresponding author: Kameswari S. Konduri. Competing Interest Statement Authors KSK and ND are listed on the patent that VGSK Technologies, Inc holds for *ProLung-budesonide*. The remaining authors declare that they have no competing interests.

## Abstract

**Background:** Inhaled budesonide benefits patients with COVID-19. *ProLung™-budesonide* enables the sustained, low dose administration of budesonide within a delivery vehicle similar to lung surfactant. *ProLung™-budesonide* may offer anti-inflammatory and protective effects to the lung in COVID-19, yet it’s effect on SARS-CoV-2 replication is unknown.

**Objective:** To determine the efficacy of *ProLung™-budesonide* against SARS-CoV-2 infection in vitro, evaluate its ability to decrease inflammation, and airway hyperresponsiveness in an animal model of lung inflammation.

**Methods:** SARS-CoV-2-infected Vero 76 cells were treated with *ProLung™-budesonide* ([0.03– 100 μg/ml]) for 3 days, and virus yield in the supernatant was measured. Ovalbumin-sensitized C57BL/6 mice received aerosolized (a) *ProLung™-budesonide* weekly, (b) only budesonide, either daily or weekly, or (c) weekly empty *ProLung™-carrier* (without budesonide). All treatment groups were compared to sensitized untreated, or normal mice using histopathologic examination, electron microscopy (EM), airway hyperresponsiveness (AHR) to Methacholine (Mch) challenge, and eosinophil peroxidase activity (EPO) measurements in bronchioalveolar lavage (BAL).

**Results:** *ProLung™-budesonide* showed significant inhibition on viral replication of SARS-CoV-2-infected cells with the selectivity index (SI) value > 24. Weekly *ProLung™-budesonide* and daily budesonide therapy significantly decreased lung inflammation and EPO in BAL. *ProLung™-budesonide* localized in type II pneumocytes, and was the only group to significantly decrease AHR, and EPO in BAL with Mch challenge

**Conclusions:** *ProLung™-budesonide* significantly inhibited viral replication in SARS-CoV-2 infected cells. It localized into type II pneumocytes, decreased lung inflammation, AHR and EPO activity with Mch challenge. This novel drug formulation may offer a potential inhalational treatment for COVID-19.

## INTRODUCTION

COVID-19 can cause significant respiratory symptoms with pulmonary compromise due to severe inflammation, often requiring ventilatory support. This process can increase airway hyperresponsiveness and possibly lead to permanent lung damage. COVID-19 can result in elevated IL-6 levels, antiphospholipid antibodies, D-dimer levels, renal failure, and increased clotting issues. (1,2).

The mechanism of COVID-19 has been shown to be secondary to SARS-CoV-2 virus binding to the ACE2 receptor on type II pneumocytes in the lung (3,4) which subsequently can result in overwhelming inflammation. Dexamethasone, a steroid, offers a significant benefit to decreasing inflammation with severe respiratory distress in COVID-19(5). Inhaled steroids such as budesonide, are also showing a decrease in the respiratory symptoms with COVID-19 (6). Other studies have shown that inhaled steroids may decrease the ACE2 receptor, which may also be beneficial in decreasing the binding of SARS-CoV-2 virus (7). While offering a significant benefit in decreasing inflammation, it is not known what effect steroids have on SARS-CoV-2 viral replication.

*ProLung™-budesonide*, uses a vehicle similar to lung surfactant, allowing for inhalational administration of a low dose of budesonide, in a sustained manner. We have previously shown in experimental animal studies, that weekly inhalation of *ProLung™-budesonide* significantly reduces lung inflammation (8). The unique lipid composition of *ProLung™-budesonide* has been shown to have immunomodulating effects, stabilize the endothelium, decrease IL-6 levels, and antiphospholipid antibodies, all of which may play an important role in COVID-19 (9–12). Studies have also shown that lung surfactant can have a protective role against SARS-CoV-2 infection (10). Our objective was to evaluate the effects of *ProLung™-budesonide* on viral replication in SARS-CoV-2 infected Vero 76 cells, and AHR with lung inflammation in ovalbumin murine model of inflammation. Electron microscopy was used to determine the stability and deposition of *ProLung™-budesonide* in the lung tissues.

## RESULTS

### Virus Yield Reduction/Neutral Red Toxicity-VYR Assay

*ProLung™-budesonide* showed highly significant antiviral activity against SARS-CoV-2, as indicated by testing with the Virus Yield Reduction)/Neutral Red Toxicity assay (**Table 3**). The EC_90_ (compound concentration that reduces viral replication by 90%) of *ProLung™-budesonide* was 4.1 μg/mL, compared to 8.1 μg/mL for the control protease inhibitor. Selectivity Index (SI_90_) was calculated as concentrations CC_50_ (50% cytotoxic, cell-inhibitory) / EC_90_ (compound concentration that reduces viral replication by 90%), by regression analysis with a SI value ≥10 considered as active. *ProLung™-budesonide* SI_90_ was >24, and for the control SI_90_ was >12.

### Airway Hyperresponsiveness (AHR) to Methacholine (Mch) Challenge

The baseline airway resistance (R_L_) in normal mice before challenge with Mch was 1.14 cm H_2_0 ml^−1^ s (**Figure 1**). The baseline R_L_ was greater in the Empty *ProLung™ carrier* and Daily budesonide treatment groups. At a cumulative dose of l mg Mch, R_L_ was increased in all groups. At the l mg Mch dose, there was no significant difference between the airway responsiveness of any of the groups of sensitized mice receiving treatment compared to the Sensitized, Untreated group. All the treatment groups except the *ProLung™-budesonide* treatment group, demonstrated a significant increase in R_L_ compared to the Normal group at a cumulative dose of 3 mg of Mch. There was no significant difference in R_L_ between the Normal Unsensitized, Untreated group and the *ProLung™-budesonide* treatment group and these were the only two groups with an R_L_ significantly less than the Sensitized, Untreated group.

**Figure 1:**
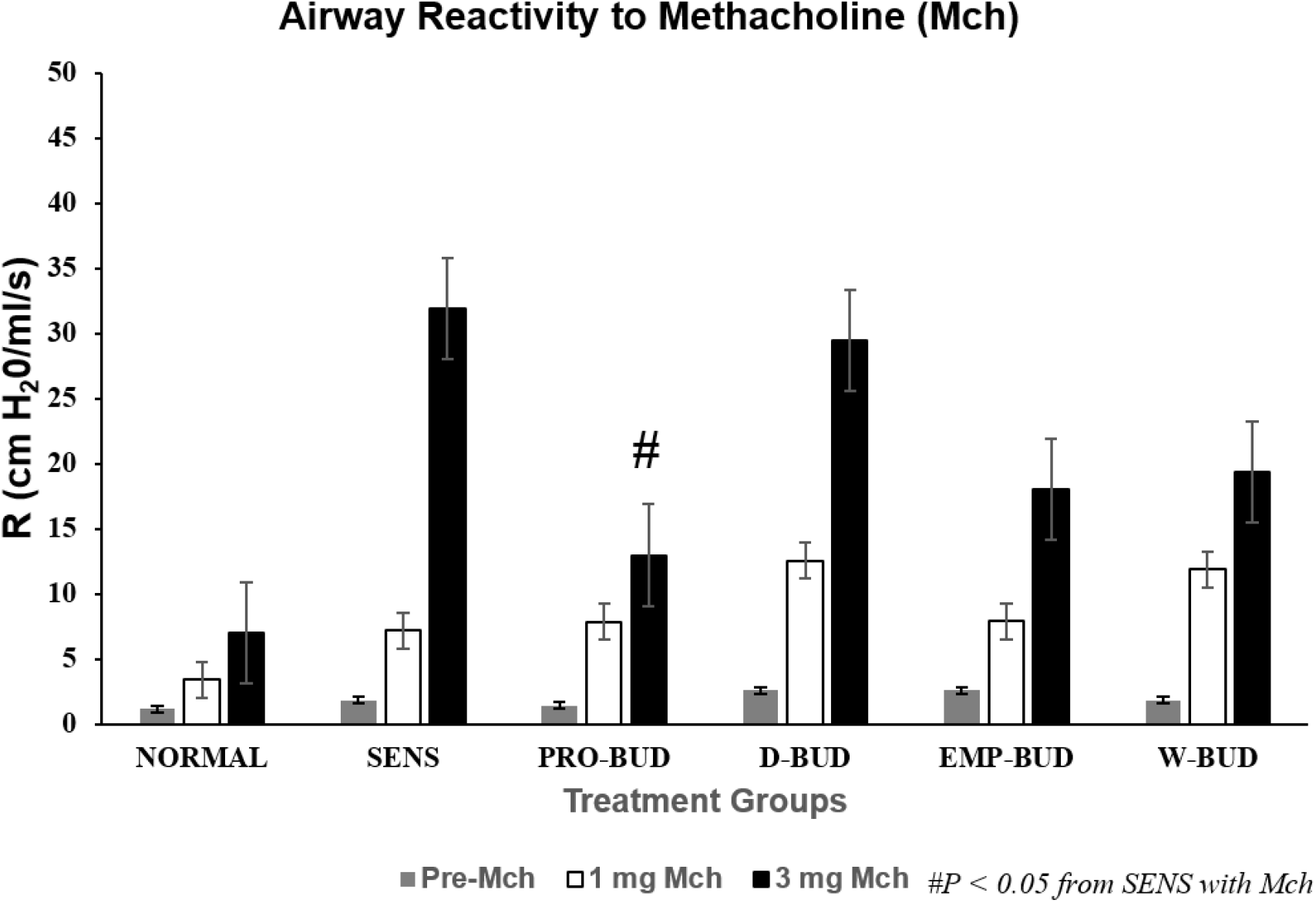
Airway Reactivity (AHR) to Methacholine (Mch) Challenge. **<u>Treatment Groups:</u>** **D-BUD =** Treatment with 20 μg of budesonide only, administered daily to sensitized mice with inflammation **PRO-BUD=**20 μg of budesonide in the ProLung™ carrier, (ProLung™-budesonide) administered as one dose, once a week to sensitized mice with inflammation **EMP-PRO=**Treatment with Empty buffer-loaded, ProLung™ carrier, administered once a week to sensitized mice with inflammation **W-BUD=**Treatment with budesonide only, administered once a week to sensitized mice with inflammation **<u>Control Groups:</u>** **NORMAL=**Normal, Untreated, Unsensitized mice **SENS=** Sensitized, Untreated, mice with inflammation. □ No methacholine challenge (NO Mch) ■ With methacholine challenge (With Mch) Airway Reactivity (AHR) to Methacholine (Mch) Challenge was measured as resistance (R in cm H_2_0/ml/s). Data is shown for baseline which is no Mch challenge (gray bar, with 1mg Mch challenge (white bar), and 3mg Mch challenge (dark bar). The baseline R_L_ was greater in the Empty *ProLung™ carrier* (EMP-PRO) and Daily budesonide (D-BUD) treatment groups. At a cumulative dose of l mg Mch, R_L_ was increased in all groups. At the l mg Mch dose, there was no significant difference between the airway responsiveness of any of the groups of sensitized mice receiving treatment compared to the Sensitized, Untreated (SENS) group. All the treatment groups except the *ProLung™-budesonide* (PRO-BUD) treatment group, demonstrated a significant increase in R_L_ compared to the Normal Unsensitized, Untreated (NORMAL) group at a cumulative dose of 3 mg of Mch. There was no significant difference in R_L_ between the Normal Unsensitized, Untreated (NORMAL) group and the *ProLung™-budesonide* (PRO-BUD) treatment groups and there were the only groups with an R_L_ significantly less than the Sensitized, Untreated (SENS) group.

### Eosinophil Peroxidase (EPO) Activity With and Without Methacholine (Mch) Challenge

In the groups without Mch challenge the *ProLung™-budesonide* (*P* < 0.001) and the Daily budesonide (*P* < 0.001) treatment groups significantly decreased the Eosinophil Peroxidase (EPO) activity in the bronchioalveolar lavage fluid (BAL), when compared to the Sensitized, Untreated group (**Figure 2**). Weekly budesonide (*P* = 0.419) and the Empty *ProLung™ carrier* (*P* = 0.213) treatment groups did not show a significant decrease in EPO activity.

**Figure 2:**
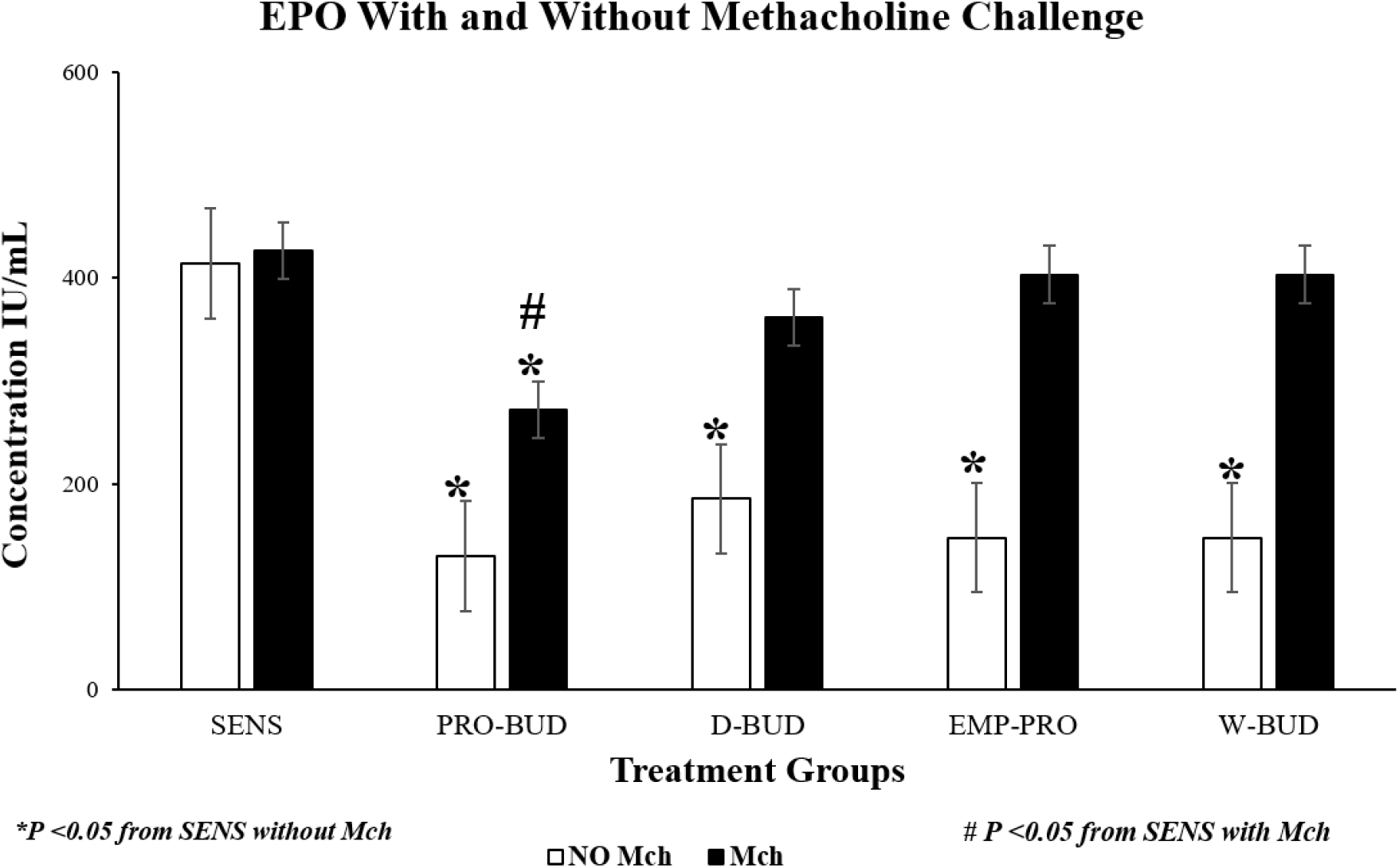
Eosinophilic Peroxidase Activity (EPO) With and Without Methacholine (Mch) Challenge. **<u>Treatment Groups:</u>** **D-BUD =** Treatment with 20 μg of budesonide only, administered daily to sensitized mice with inflammation **PRO-BUD=**20 μg of budesonide in the ProLung™ carrier, (ProLung™-budesonide) administered as one dose, once a week to sensitized mice with inflammation **EMP-PRO=**Treatment with Empty buffer-loaded, ProLung™ carrier, administered once a week to sensitized mice with inflammation **W-BUD=**Treatment with budesonide only, administered once a week to sensitized mice with inflammation **<u>Control Groups:</u>** **NORMAL=**Normal, Untreated, Unsensitized mice **SENS=** Sensitized, Untreated, mice with inflammation. □ No methacholine challenge (NO Mch) ■ With methacholine challenge (With Mch) Graph represents cumulative results from a 4-week study of eosinophilic peroxidase activity (EPO), a marker of inflammation, measured in bronchoalveolar lavage fluid (BAL) and Airway Reactivity (AHR) to Methacholine (Mch) Challenge. In the groups Without Mch(NO Mch) challenge all the treatment groups showed a significant decrease in EPO activity, when compared to the Sensitized, Untreated (SENS) group. Only the weekly treatments with *ProLung™-budesonide* (PRO-BUD) significantly decreased EPO activity, with Mch and without Mch (NO Mch) challenge when compared to the Sensitized, Untreated (SENS) group. Daily budesonide (D-BUD), Weekly budesonide (WK-BUD and the Empty *ProLung™ carrier* (EMP-PRO) treatment groups did not show a significant decrease in EPO activity with Mch challenge.

With Mch challenge, EPO activity of the all groups was increased, except for the *ProLung™-budesonide* treated group, which showed a significant decrease in EPO activity *P* < 0.005). There was no significant difference in the EPO activity, with or without Mch challenge, only in the *ProLung™-budesonide* treated (*P* = 0.68) and the Normal Unsensitized, Untreated group. Normal Unsensitized, Untreated group had no detectable EPO activity in the BAL.

### Lung Histology

Examples of lung tissues from the treatment groups are shown in **Figures 3 and 4** (100x magnification, hematoxylin-eosin). The lung tissues from the Sensitized, Untreated (SENS) mice had persistent and significant inflammation, including accumulation of inflammatory cells in bronchiolar, peribronchiolar, and perivascular tissues, along with significant submucosal thickening and epithelial hyperplasia, during the 4-week period. Lung inflammation was markedly increased along with bronchoconstriction, cellular infiltrates with methacholine (With Mch) challenge in all the groups except for the Normal Unsensitized, Untreated and *ProLung™-budesonide* treatment group. *ProLung™-budesonide* was the only treatment group that did not show a significant increase in lung inflammation, with (With Mch) or without Mch (NO Mch) challenge, when compared to the Sensitized, Untreated group. Daily budesonide treatment group only showed reduction in lung inflammation without Mch challenge. The daily budesonide group treatment group showed marked increase in inflammation along with bronchoconstriction and cellular infiltrates with Mch challenge.

**Figure 3:**
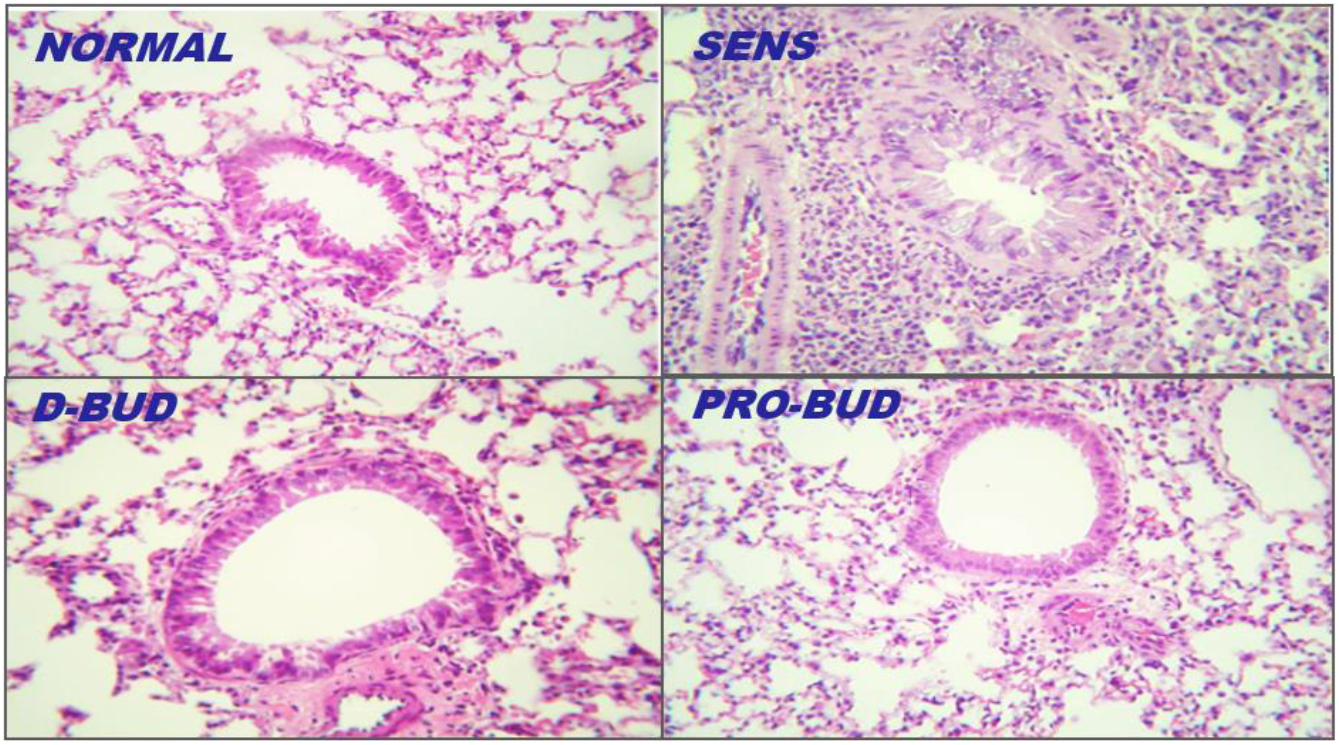
Lung Histology, with and without methacholine (Mch) challenge. **<u>Treatment Groups:</u>** **D-BUD =** Treatment with 20 μg of budesonide only, administered daily to sensitized mice with inflammation **PRO-BUD=**20 μg of budesonide in the ProLung™ carrier, (ProLung™-budesonide) administered as one dose, once a week to sensitized mice with inflammation **EMP-PRO=**Treatment with Empty buffer-loaded, ProLung™ carrier, administered once a week to sensitized mice with inflammation **W-BUD=**Treatment with budesonide only, administered once a week to sensitized mice with inflammation **<u>Control Groups:</u>** **NORMAL=**Normal, Untreated, Unsensitized mice **SENS=** Sensitized, Untreated, mice with inflammation. Examples of lung tissues from the treatment groups are shown in **Figures 3 and 4** (100x magnification, hematoxylin-eosin). The lung tissues from the Sensitized, Untreated (SENS) mice had persistent and significant inflammation, including accumulation of inflammatory cells in bronchiolar, peribronchiolar, and perivascular tissues, along with significant submucosal thickening and epithelial hyperplasia, during the 4-week period. Lung inflammation was markedly increased along with bronchoconstriction, cellular infiltrates with methacholine (With Mch) challenge in all the groups except for the NORMAL and *ProLung™-budesonide* (PRO-BUD) treatment groups. *ProLung™-budesonide* (PRO-BUD) was the only treatment group that did not show a significant increase in lung inflammation, with (With Mch) or without Mch (NO Mch) challenge, when compared to the Sensitized, Untreated (SENS) group. Daily budesonide treatment (D-BUD) group only showed reduction in lung inflammation without Mch challenge. The daily budesonide group (D-BUD) group showed marked increase in inflammation along with bronchoconstriction and cellular infiltrates with Mch challenge.

**Figure 4:**
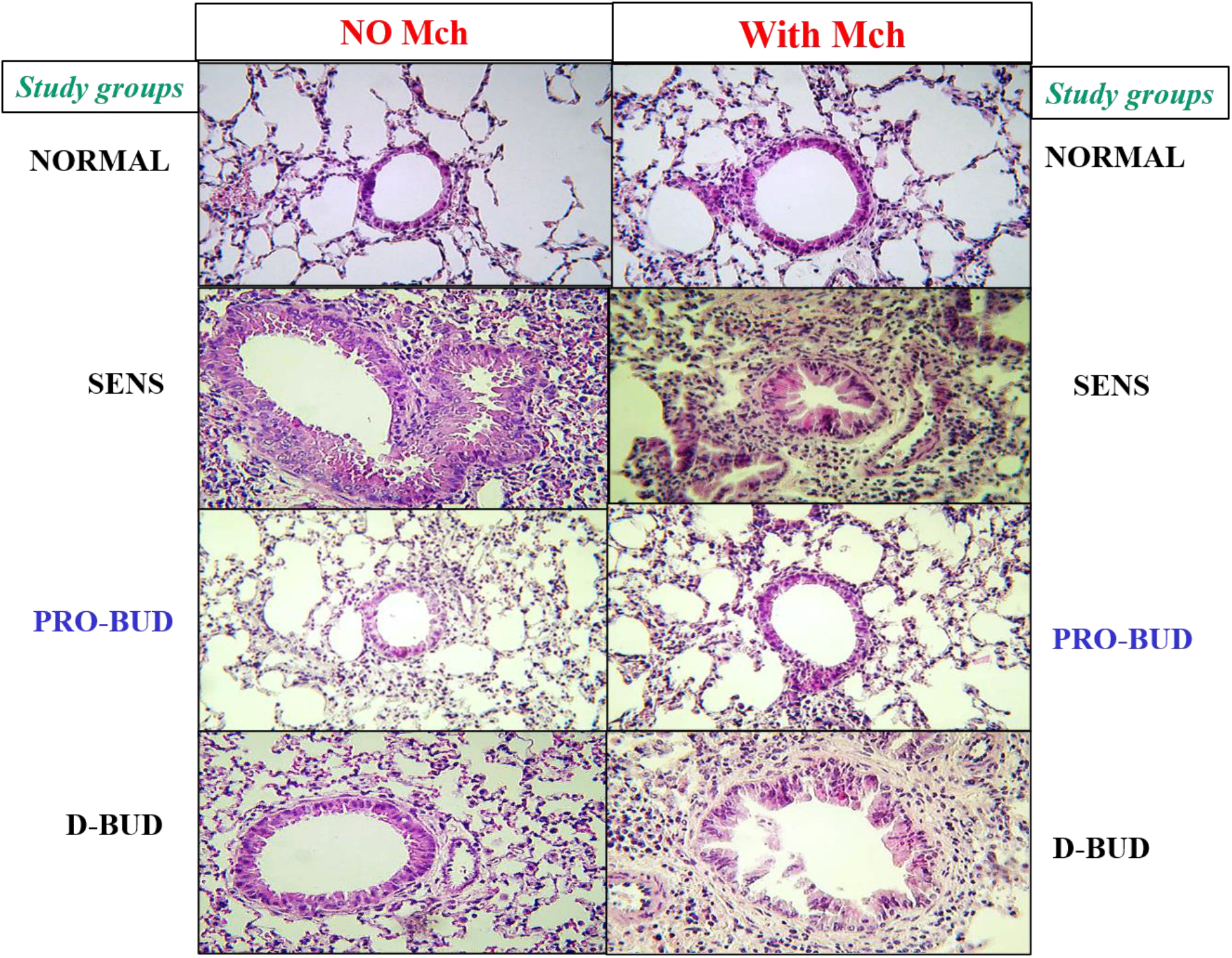
Lung Histology, with and without methacholine (Mch) challenge. **<u>Treatment Groups:</u>** **D-BUD =** Treatment with 20 μg of budesonide only, administered daily to sensitized mice with inflammation **PRO-BUD=**20 μg of budesonide in the ProLung™ carrier, (ProLung™-budesonide) administered as one dose, once a week to sensitized mice with inflammation **EMP-PRO=**Treatment with Empty buffer-loaded, ProLung™ carrier, administered once a week to sensitized mice with inflammation **W-BUD=**Treatment with budesonide only, administered once a week to sensitized mice with inflammation **<u>Control Groups:</u>** **NORMAL=**Normal, Untreated, Unsensitized mice **SENS=** Sensitized, Untreated, mice with inflammation. Examples of lung tissues from the treatment groups are shown in **Figures 3 and 4** (100x magnification, hematoxylin-eosin). The lung tissues from the Sensitized, Untreated (SENS) mice had persistent and significant inflammation, including accumulation of inflammatory cells in bronchiolar, peribronchiolar, and perivascular tissues, along with significant submucosal thickening and epithelial hyperplasia, during the 4-week period. Lung inflammation was markedly increased along with bronchoconstriction, cellular infiltrates with methacholine (With Mch) challenge in all the groups except for the NORMAL and *ProLung™-budesonide* (PRO-BUD) treatment groups. *ProLung™-budesonide* (PRO-BUD) was the only treatment group that did not show a significant increase in lung inflammation, with (With Mch) or without Mch (NO Mch) challenge, when compared to the Sensitized, Untreated (SENS) group. Daily budesonide treatment (D-BUD) group only showed reduction in lung inflammation without Mch challenge. The daily budesonide group (D-BUD) group showed marked increase in inflammation along with bronchoconstriction and cellular infiltrates with Mch challenge.

### Histopathology Score With and Without Methacholine (Mch) Challenge

The lung tissues from the Sensitized, Untreated group had persistent and significant inflammation, without Methacholine (Mch) challenge, including accumulation of inflammatory cells in bronchiolar, peribronchiolar, perivascular tissues, and alveolar regions along with significant submucosal thickening and epithelial hyperplasia, during the four-week period (**Figure 5**). The inflammation was markedly increased with bronchoconstriction and cellular infiltrates with Mch challenge.

**Figure 5:**
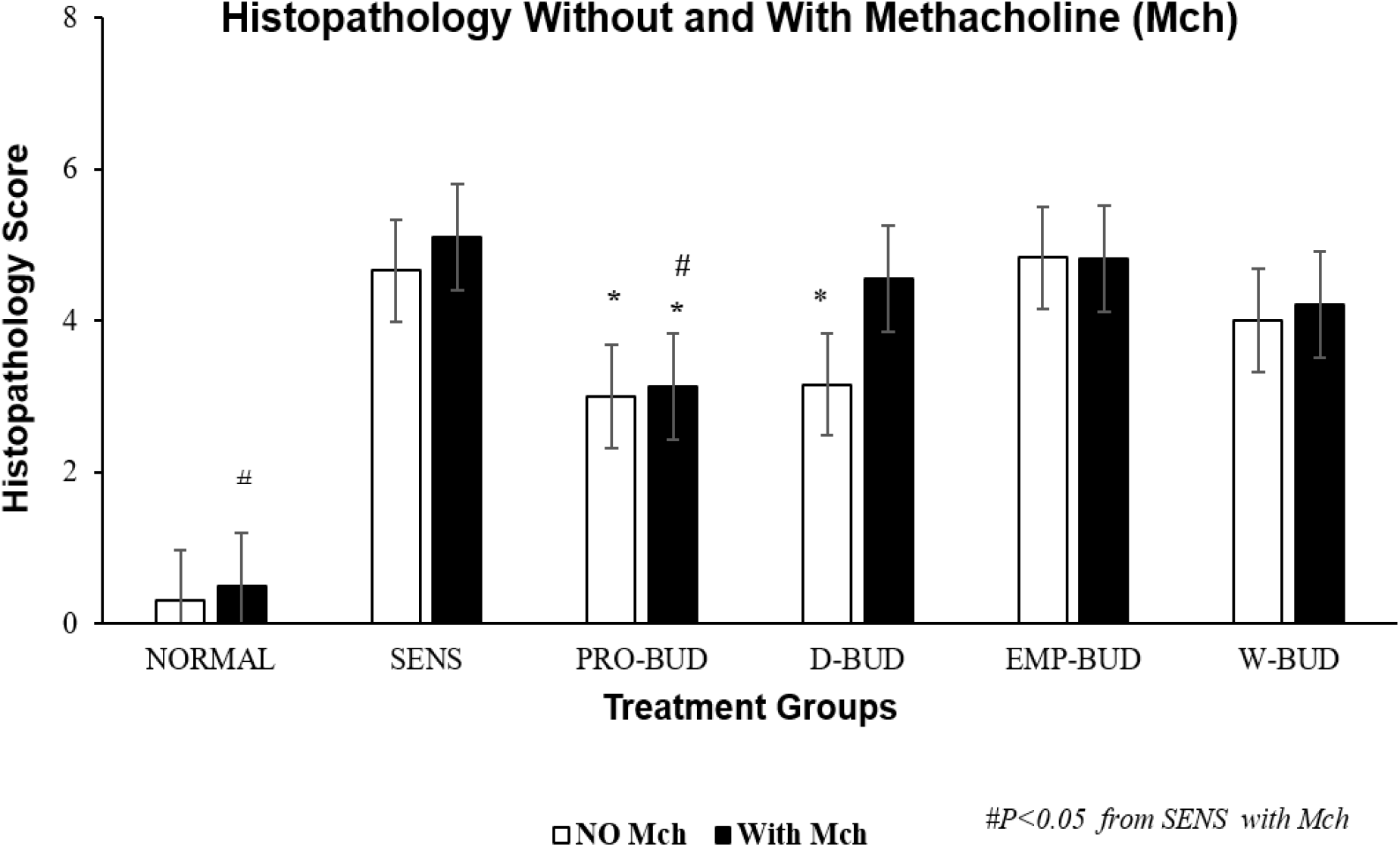
Lung Histopathology Scores With and Without Methacholine (Mch) Challenge. **<u>Treatment Groups:</u>** **D-BUD =** Treatment with 20 μg of budesonide only, administered daily to sensitized mice with inflammation **PRO-BUD=**20 μg of budesonide in the ProLung™ carrier, (ProLung™-budesonide) administered as one dose, once a week to sensitized mice with inflammation **EMP-PRO=**Treatment with Empty buffer-loaded, ProLung™ carrier, administered once a week to sensitized mice with inflammation **W-BUD=**Treatment with budesonide only, administered once a week to sensitized mice with inflammation **<u>Control Groups:</u>** **NORMAL=**Normal, Untreated, Unsensitized mice **SENS=** Sensitized, Untreated, mice with inflammation. □ No methacholine challenge (NO Mch) ■ With methacholine challenge (With Mch) Graph depicts the cumulative histopathology score from a 4-week study, with and without methacholine (Mch) challenge. Scores were obtained from a scoring system (**Table 1**) and were determined by a veterinary pathologist blinded to the treatment groups. The lung tissues from the Sensitized, Untreated (SENS) group had persistent and significant inflammation, without methacholine (No Mch) challenge which was increased with methacholine challenge (With Mch). There was a significant reduction in total lung histopathology score without Mch challenge, in the *ProLung™-budesonide* (PRO-BUD) and Daily budesonide (D-BUD) treatment groups when compared to the Sensitized, Untreated (SENS) group. Similar decreases were not observed with the other treatment groups. Only the *ProLung™-budesonide* (PRO-BUD) treatment group with the Mch challenge, had a significant decrease in total histopathology score when compared to the Sensitized, Untreated (SENS) group. There was also a significant decrease in lung inflammation in the *ProLung™-budesonide* (PRO-BUD) group in comparison with the Weekly budesonide (WK-BUD) group. None of the other treatment groups, including the Daily budesonide (D-BUD) treatment group, Weekly budesonide (WK-BUD), or Empty *ProLung™ carrier* (EMP-PRO) treatment groups showed a similar reduction with Mch challenge.

There was a significant reduction in total lung histopathology score without Mch challenge, in the *ProLung™-budesonide* (*P* < 0.020) and Daily budesonide (*P <* 0.030) treatment groups when compared to the Sensitized, Untreated group. Similar decreases were not observed with the other treatment groups. Only the *ProLung™-budesonide* treatment group with Mch challenge, had a significant decrease in total histopathology score (*P* < 0.0009) when compared to the Sensitized, Untreated group. None of the other treatment groups (including Daily budesonide treatment group) did not show a similar reduction with Mch challenge.

### *ProLung™-budesonide* Localizes to Type II pneumocytes in the Lung

Scanning electron microscopy showed the deposition of the *ProLung™-budesonide* in the lung a week after a single dose was administered **(Figure 6**). Results show that *ProLung™* -*budesonide* was taken up into Type II pneumocytes at the alveolar level in the lung tissues. *ProLung™-budesonide* was detected upto 10 days post dosing, and was not detected at the two-week period after a single dose was administered.

**Figure 6:**
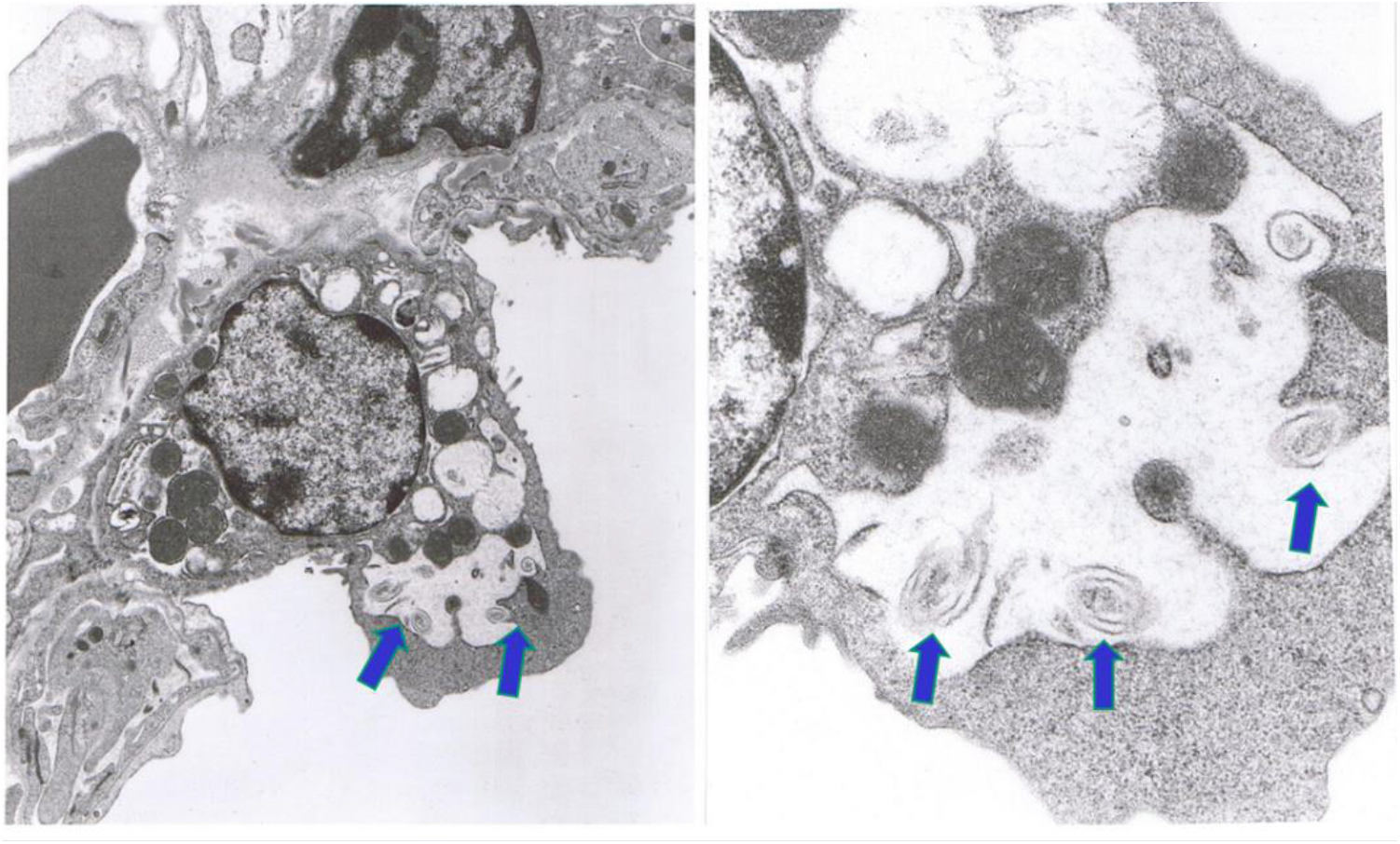
ProLung™budesonide Localizes to Type II pneumocytes in the Lung. Scanning electron microscopy showed the deposition of *ProLung™-budesonide* in the lung after a week after a single dose was administered. Arrows depict the swirls inside the Type II pneumocytes. Top left lower magnification and top right higher magnification. *ProLung™*-*budesonide* was taken up into the Type II pneumocytes at the alveolar level in the lung tissues.

## DISCUSSION

COVID-19 can cause significant respiratory symptoms with compromise that may require ventilatory support, with prolonged lung inflammation. Pulmonary inflammation can lead to lung damage and an increase in airway hyperreactivity (AHR), which can subsequently result in the respiratory symptoms and compromise. After exposure, SARS-CoV-2 infection is thought to occur through binding to the ACE2 receptor on type II pneumocytes in the lung, and the gastrointestinal (GI) mucosa (3,4) which subsequently can result in overwhelming inflammation.

Corticosteroids have demonstrated anti-inflammatory effects in COVID-19 patients (5). Dexamethasone, offers a significant benefit in decreasing inflammation in COVID-19 patients with severe respiratory distress. Inhaled steroid, budesonide, has recently been shown to decrease respiratory symptoms. and possible progression to COVID-19 (6). Other studies have shown that inhaled steroids may decrease the ACE2 receptor, which may also be beneficial in decreasing SARS-CoV-2 binding (7). While offering a significant benefit in decreasing inflammation, it is not known what effect steroids have on viral replication. Lung surfactant may have a protective role against SARS-CoV-2 infection (10). *ProLung™ budesonide,* is similar in composition to lung surfactant which allows for weekly administration of budesonide in a sustained carrier.

In this study, *ProLung™ budesonide* significantly reduced viral replication of SARS-CoV-2 in *Vero* cells (**Table 3**), airway hyperresponsiveness (AHR) to methacholine (Mch) challenge (**Figure 1**), and lung inflammation (**Figures 3-*5***). *ProLung™ budesonide,* also showed a significant decrease in eosinophil peroxidase activity (EPO) a marker of inflammation, and AHR with Mch challenge (**Figure 2**). Daily therapy with budesonide, decreased lung inflammation, but did not show a prolonged effect, or have an effect on decreasing AHR or EPO activity with Mch challenge (**Figures 1-5**).

We noted on electron microscopy studies that *ProLung™ budesonide* localizes in type II pneumocytes (**Figure 6**), the site of SARS-CoV-2 binding. Type II pneumocytes produce and secrete pulmonary surfactant lipids and proteins, and other soluble components of the innate immune system (10–12). They are considered to be the regulatory cells of the lung, may play a critical role in lung inflammation, with immune interactions with alveolar macrophages (11,16–18). In diseases such as tuberculosis (TB), Type II pneumocytes likely signal alveolar macrophages to retain the TB organism subsequently leading to lung inflammation. Studies have shown that lung macrophages also play an important role in the lung inflammation and damage in COVID-19 and acute respiratory distress syndrome (ARDS) (11,16–19).

*ProLung™ budesonide* is delivered to the alveolar junction, targets Type II pneumocytes, and is noted to decrease SARS-CoV-2 viral replication. Dr. PRJ Gangadharam (VGSK founding scientist) and Dr. Düzgüneş have performed extensive work on antibiotics encapsulated in the *ProLung™-carrier* and noted the targeting to the macrophages of the reticuloendothelial system (14, 20, 21). They also noted the *ProLung™-carrier* preferentially target areas with increased inflammation and macrophages (14,20, 21) in the systemic circulation.

*ProLung™ budesonide* may have a potential to interrupt the interaction of Type II and alveolar macrophages, and the subsequent progression to the lung inflammation noted in COVID-19. In addition, it may aid in lung stabilization and maintaining alveolar function, secondary it’s sustained steroid effect with a composition similar to lung surfactant. The unique lipid composition of *ProLung™ budesonide*, may play a role in the innate immune system and may decrease IL-6 levels which can be markedly elevated in COVID-19 (1, 2). With these unique properties, *ProLung™ system* can have a significant impact in treating the COVID-19 Pandemic.

Pegylated delivery systems are now being used to deliver a variety of immune based therapies and mRNA vaccines, such as Moderna® and Pfizer® (22–24) and have been implicated in allergic reactions. Similarly, Poly ethylene glycol (PEG) 2000 is also used in *ProLung™ budesonide* but have not encountered similar allergic reactions. To determine the safety of *ProLung™ budesonide*, we have conducted long-term, safety, and toxicity studies in an allergic model, using a repeat allergen challenge. We noted significant decreases in markers of allergic inflammation such as serum IgE levels, reduction of eosinophils in the lung lavage fluid and peripheral blood, and EPO activity (8, 15). There were no issues noted with toxicity or severe allergic reactions during dose escalation and long-term toxicity studies (unpublished data) conducted in our animal studies.

*ProLung™ budesonide* is delivered as one dose, weekly as an inhalation, and has many differences in composition from the vaccine delivery systems delivered as intramuscular injections. PEG 2000 also has an immunomodulatory function (23–25). The small amount of PEG in our carrier system may also act as an additional barrier to prevent viral attachment of SARS-CoV-2 virus. COVID-19 may result in the production of autoimmune antibodies, such as antiphospholipid antibodies, which may be directed at lung surfactant lipids (9) *ProLung™-budesonide,* secondary to it’s surfactant like composition, may be crucial in treating COVID-19 respiratory symptoms as well post COVID-19 syndrome, such as the “long haulers” who have lung symptoms for months post the initial infection (26). In addition, it has a stabilizing effect on the lung mucosa which may aid in decreasing the fibrotic changes in the lung (unpublished Data). In addition to the anti-inflammatory effect, the *ProLung™-budesonide* decreases AHR to Mch challenge without the addition of a beta agonist, and may decrease airway remodeling, which is not noted with daily budesonide therapy.

*ProLung™-budesonide* may improve patient compliance as it offers a less frequent dosing for chronic respiratory diseases, and possibly for the “long haulers” post COVID-19. Daily dosing of a medication may lead to problems of noncompliance and treatment failures, which might result in increased hospitalizations and complications. *ProLung™-budesonide* offers a therapy that can be administered in a safe, effective manner as an inhalation, with a low dose of steroid in a carrier similar to composition to surfactant targeted in the lung to the point of viral attachment of SARS-CoV-2. With these unique properties, *ProLung™-budesonide* can have a significant impact in treating the COVID-19 Pandemic.

## MATERIALS AND METHODS

### Virus Yield Reduction (VYR) Assay

A Virus Yield Reduction (VYR) assay was performed to determine test compound inhibition of virus replication. Confluent or near-confluent cell culture monolayers of Vero 76 cells were prepared in 96-well microplates. *ProLung™-budesonide* was tested at eight half-log_10_ concentrations (0.032, 0.1, 0.32, 1, 3.2, 10, 32 and 100 μg/ml) for antiviral activity and cytotoxicity. Plates were incubated at 37 °C with 5% CO_2_ until >80% CPE (virus-induced cytopathic effect) was observed in virus control wells. Five microwells were used per dilution: three for infected cultures and two for uninfected toxicity cultures. Controls for the experiment consisted of six microwells that were infected and not treated (virus controls) and six that were untreated and uninfected (cell controls) on every plate. A known active drug was tested (protease inhibitor) in parallel as a positive control drug. Cells were scored for the presence or absence of virus after distinct CPE was observed, and the CCID_50_ (50% cell culture infectious dose) is calculated using the Reed-Muench method (13). In addition, virus yielded in the presence of *ProLung™-budesonide* was titrated and compared to virus titers from the untreated virus controls. Titration of the viral samples was performed by endpoint dilution.

After maximum virus-induced cytopathic effect (CPE) was observed, the viable plates were stained with 0.011% neutral red dye at 37 °C. The neutral red medium was removed, and the cells rinsed once with phosphate buffered saline (PBS) to remove residual dye. The incorporated dye content was extracted and quantified by evaluation of absorbance on a spectrophotometer at 540 nm. The dye content in each set of wells was converted to a percentage of dye present in untreated control wells and normalized based on the virus control. The 90% (one log_10_) effective concentration (EC_90_) was calculated by regression analysis by plotting the log_10_ of the inhibitor concentration versus log_10_ of virus produced at each concentration. The 50% effective (EC_50_, virus-inhibitory) concentrations and 50% cytotoxic (CC_50_, cell-inhibitory) concentrations were then calculated by regression analysis. The quotient of CC_50_ divided by EC_50_ gives the selectivity index (SI) value, with compounds having a SI value ≥10 being considered active.

### Animal Studies

Six-week-old male C57 black 6 mice (C57BL/6) were purchased from Charles River Laboratories, Inc., Wilmington, MA. The animals were provided with an ovalbumin-free diet and water *ad libitum* and were housed in an environment-controlled, pathogen-free animal facility. All animal protocols were approved by the Animal Care Committee of the Medical College of Wisconsin and the Zablocki Veterans Administration Medical Center, in agreement with the National Institute of Health’s guidelines for the care and use of laboratory animals. We were unable to conduct our studies in animal models with COVID-19 as current animal models generally had only mild forms of the disease and were not seen as adequate models for assessment of anti-inflammatory properties of inhalation drugs at the time our studies were conducted.

### Sensitization

The C57BL/6 mice were sensitized with ovalbumin (OVA) as described in our previous studies (8). This method of sensitization led to a significant elevation in eosinophil peroxidase (EPO) levels in the bronchioalveolar lavage fluid (BAL), and lung inflammation by day 24, as seen by histopathology. This method also increased airway hyperresponsiveness (AHR) to methacholine (Mch) challenge, by day 24. All treatment groups were compared with either Sensitized, Untreated or Normal, Unsensitized, Untreated mice.

The dose of budesonide was extrapolated from our previous dose-response studies (8). The 20 μg dose of budesonide was noted to decrease EPO in the BAL, and inflammation on histopathological examination of the lung tissues, along with other inflammatory parameters studied, without evidence of toxicity to the spleen, liver, bone marrow, skin or the gastrointestinal tract. Based on our results, 20 μg of budesonide was encapsulated as *ProLung™-budesonide* for administration one dose, once a week of as an inhalation.

### Study Groups

Therapy was initiated on day 25, one day after the OVA sensitization was completed. Sensitized animals received nebulized treatments for four weeks. Each study group consisted of 20 mice and was followed for a four-week period. Five animals from each treatment group and from each of the two control groups, sensitized and unsensitized, were euthanized by means of an overdose of methoxyflurane inhalation, 24 hours after the first treatments were given, and then at weekly intervals for four weeks. At each time point, measurements of EPO in BAL were obtained and histopathologic examination of the lung tissues was performed.

### Treatment Groups

After the OVA sensitization was completed (day 25), Sensitized animals received nebulized treatments for four weeks as follows (**Table 1**): (a) (PRO-BUD)-received 20 μg of *ProLung™-budesonide* administered once a week; (b) (D-BUD)-20 μg of budesonide (without *ProLung™ carrier*) administered daily (c) (EMP-PRO)-received Empty *ProLung™ carrier* (buffer-loaded), administered once a week; (d) (W-BUD)-20 μg of budesonide (without *ProLung™*) administered once a week. All treatment groups were compared to either Sensitized Untreated (SENS) or Untreated, Unsensitized (NORMAL) mice.

**Table 1:** Study Groups. **<u>Treatment Groups:</u>** **D-BUD =** Treatment with 20 μg of budesonide only, administered daily to sensitized mice with inflammation **PRO-BUD=**20 μg of budesonide in the ProLung™ carrier, (ProLung™-budesonide) administered as one dose, once a week to sensitized mice with inflammation **EMP-PRO=**Treatment with Empty buffer-loaded, ProLung™ carrier, administered once a week to sensitized mice with inflammation **W-BUD=**Treatment with budesonide only, administered once a week to sensitized mice with inflammation **<u>Control Groups:</u>** **NORMAL=**Normal, Untreated, Unsensitized mice **SENS=** Sensitized, Untreated, mice with inflammation.

| GROUP | Treatment Type | *ProLung*<sup>TM</sup> | Budesonide | Frequency |
| --- | --- | --- | --- | --- |
| **PRO-BUD** | Weekly treatment with *ProLung*<sup>TM</sup>-budesonide | + | + | Once per week |
| **EMP-PRO** | Buffer-Loaded Empty *ProLung*<sup>TM</sup> carrier | + | - | Once per week |
| **D-BUD** | Daily treatment with budesonide only | - | + | Daily |
| **W-BUD** | Weekly treatment with budesonide only | - | + | Once per week |
| **SENS** | Sensitized Untreated with inflammation | - | - | None |
| **NORMAL** | Normal, Unsensitized Untreated | - | - | None |

### Drugs and Reagents

Budesonide for daily therapy was diluted from premixed vials (0.25 mg/ml) commercially available from Astra Pharmaceuticals (Wayne, PA), and administered via a Salter Aire Plus Compressor (Salter Labs, Irvine, CA). Budesonide for encapsulation, N-2-hydroxethylpiperzine-N’-2-ethanesulfonic acid (HEPES), ovalbumin, methacholine, PBS, sodium citrate, 0-phenylenediamine, 4N H_2_SO_4_ and horseradish peroxidase were purchased from Sigma-Aldrich, St. Louis, MO. Phosphatidylcholine (PC), phosphatidylglycerol (PG), and poly (ethylene glycol)-distearoylphosphatidylethanolamine (PEG-DSPE) were obtained from Avanti Polar Lipids, Alabaster, AL.

### *ProLung™-budesonide* Preparation

Budesonide was encapsulated into the *ProLung™ carrier* as previously described (8, 14,15). Lipids were dried onto the sides of a round-bottomed glass flask or glass tube by rotary evaporation. The dried film was then hydrated by adding sterile 140 mmol/L NaCl and 10 mmol/L HEPES (pH 7.4) and vortexing. The resulting multilamellar liposomes were extruded 11 times through two stacked polycarbonate membranes of 0.8 μm pore diameter (Whatman-Nuclepore, Sigma-Aldrich) using a custom-built high-pressure extrusion device or a syringe extruder (Avanti Polar Lipids).. Empty *ProLung™ carrier* was prepared similar to *ProLung™-budesonide*, without budesonide, and was diluted with HEPES-buffered saline to maintain an equal volume for dosing.

### Histopathology

Histopathological examinations were performed on lungs that were removed and fixed with 10% phosphate-buffered formalin as previously described (8). Tissue samples were taken from the trachea, bronchi, large and small bronchioles, interstitium, alveoli, and pulmonary blood vessels. The tissues were embedded in paraffin, sectioned at 5μm thickness and stained with hematoxylin and eosin, and analyzed using light microscopy at 100x magnification. Coded slides were examined by a veterinary pathologist in a blinded fashion for evidence of inflammatory changes. (**Table 2**). Each of the parameters evaluated was given an individual number score. Objective measurements of histopathological changes included the number of eosinophils and other inflammatory cells, surrounding the bronchi, aggregation around blood vessels, presence of desquamation and hyperplasia of the airway epithelium, mucus formation in the lumen of the airways and infiltration of inflammatory cells surrounding the alveoli. Each of the parameters evaluated were given an individual number score. The cumulative score was obtained using the individual scores and designated as no inflammation (*score:*0), mild inflammation (*score:*1–2), moderate inflammation (*score:*3–4), and severe inflammation (*score:*5–6).

**Table 2:** Quantitative Histopathology Scoring System. Histopathologic examination was performed lungs that were removed and fixed with 10% phosphate-buffered formalin. Tissue samples were taken from the trachea, bronchi, large and small bronchioles, interstitium, alveoli, and pulmonary blood vessels. The tissue slides were stained with hematoxylin and eosin, and analyzed through use of light microscopy at a magnification of 100x. Coded slides were examined by a veterinary pathologist, in a blinded fashion, for evidence of inflammatory changes, including (1) bronchiolar epithelial hyperplasia and wall thickening, (2) bronchiolar, peribronchiolar, and perivascular edema, and (3) accumulation of eosinophils, neutrophils, and mononuclear inflammatory cells. Each of the parameters evaluated was given an individual numerical score. The cumulative score was obtained through use of the individual scores; inflammation was designated as none (score, 0), mild (score, 1-2), moderate inflammation (score, 3-4), or severe inflammation (score, 5-6).

| TRACHEA | BRONCHI | LARGE BRONCHIOLES | SMALL BRONCHIOLES | ALVEOLAR INTERSTITIUM | Alveoli |
| --- | --- | --- | --- | --- | --- |
| **Epithelium**<br>Hyperplasia(mm) | **Epithelium**<br>Hyperplasia(mm) | **Epithelium**<br>Hyperplasia(mm) | **Epithelium**<br>Hyperplasia(mm) | **Thickening(mm)**<br>Edema(mm)<br>Cells(#)-PMNs(#),<br>Eosinophils(#) | **Thickening(mm)**<br>Edema(mm)<br>Cells(#)-PMNs(#),<br>Eosinophils(#)<br>Multinucleated-Giant Cells(#) |
| Desquamation | Desquamation | Desquamation | Desquamation |  |  |
| **Submucosa**<br>Edema(mm)<br>Cells(#)-PMNs(#),<br>Eosinophils(#) | **Submucosa**<br>Edema(mm)<br>Cells(#)-PMNs(#),<br>Eosinophils(#) | **Submucosa**<br>Edema(mm)<br>Cells(#)-PMNs(#)<br>Eosinophils(#) | **Submucosa**<br>Edema(mm)<br>Cells(#)-PMNs(#)<br>Eosinophils(#) | **Microgranulomas**<br>Cells(#)-PMNs(#),<br>Eosinophils(#)<br>Multinucleated-Giant Cells(#) |  |
| **Granulomas** | **Granulomas** | **Granulomas** | **Granulomas** | **Microranulomas** |  |
| **Blood Vessels**<br>Perivascular edema<br>Perivascular cuffing<br>Cells(#)-PMNS(#),<br>Eosinophils(#) | **Blood Vessels**<br>Perivascular edema<br>Perivascular cuffing<br>Cells(#)-PMNS(#),<br>Eosinophils(#) | **Blood Vessels**<br>Perivascular edema<br>Perivascular cuffing<br>Cells(#)-PMNS(#),<br>Eosinophils(#) | **Blood Vessels**<br>Perivascular edema<br>Perivascular cuffing<br>Cells(#)-PMNS(#),<br>Eosinophils(#) | **Blood Vessels**<br>Perivascular edema<br>Perivascular cuffing<br>Cells(#)-PMNS(#),<br>Eosinophils(#) | **Blood Vessels**<br>Perivascular edema<br>Perivascular cuffing<br>Cells(#)-PMNS(#),<br>Eosinophils(#) |

**Table 3:** ProLung™-budesonide activity in Vero cells. *ProLung™-budesonide* showed highly significant antiviral activity against SARS-CoV-2, as indicated by testing with the Virus Yield Reduction)/Neutral Red Toxicity assay. The 50% effective (EC_50_, virus-inhibitory) concentrations and 50% cytotoxic (CC_50_, cell-inhibitory) concentrations were then calculated by regression analysis. The quotient of CC_50_ divided by EC_50_ gives the selectivity index (SI) value, with compounds having a SI value ≥10 being considered active. The EC_90_ (compound concentration that reduces viral replication by 90%) of *ProLung™-budesonide* was 4.1 μg/mL, compared to 8.1 μg/mL for the control protease inhibitor. The SI_90_ calculated as CC_50_/EC_90_ for *ProLung™-budesonide* was >24, and for the control was >12.

| Groups | *ProLung*<sup>TM</sup>-budesonide | Control-Protease inhibitor |
| --- | --- | --- |
| EC<sub>90</sub> | 4.1 | 8.1 |
| CC<sub>50</sub> | >100 | >100 |
| SI<sub>90</sub> | >24* | >12 |

### Airway Hyperresponsiveness (AHR) To Methacholine (Mch) challenge

The effectiveness of the drug and drug-carrier combination on airway hyperreactivity (AHR) to methacholine (Mch) challenge was evaluated by measuring Pulmonary Mechanics using the protocols as previous described (15). AHR was measured in spontaneously breathing tracheally intubated mice that received up to 3 mg of Mch given intraperitoneally. Pulmonary resistance measurements were made after four weeks of therapy. As an antigen challenge and to demonstrate sensitization, an aerosolized dose of 6% ovalbumin was given to each animal 24 hours before the evaluation of the pulmonary mechanics. The digitalized Data were analyzed for dynamic pulmonary compliance, pulmonary resistance, tidal volume, respiratory frequency and minute ventilation from six to ten consecutive breaths at each recording event. Compliance and resistance were calculated from pleural pressure, airflow, and volume data. Mch challenge was performed after baseline measurements were obtained. Mch was injected intraperitoneally at three-minute intervals in successive cumulative doses of 30, 100, 300, 1,000 and 3,000μg.

### Eosinophil Peroxidase (EPO) Activity in Bronchoalveolar Lavage (BAL) Fluid

EPO activity was measured in the BAL, with and without methacholine (Mch) challenge. At the time of sacrifice, the trachea was exposed and cannulated with a ball-tipped 24-gauge needle. The lungs were lavaged three times with 1 ml phosphate-buffered saline (PBS). All washings were pooled and the samples were frozen at −70°C. The samples were later thawed and assayed to determine EPO activity. A substrate solution consisting of 0.1 mol/L sodium citrate, 0-phenylenediamine, and H_2_O_2_(3%), pH 4.5, was mixed with BAL supernatants at a ratio of 1:1. The reaction mixture was incubated at 37°C, and the reaction was stopped by the addition of 4 N H_2_SO_4_. Horseradish peroxidase was used as a standard. EPO activity (in international units per milliliter) was measured by spectrophotometric analysis at 490 nm.

### Electron Microscopy

Lung specimens were processed using standard protocols and were evaluated under transmission electron microscopy to evaluate using a Hitachi 600 electron microscope. Data was evaluated to determine the stability and deposition of the *ProLung™-budesonide* in the lung. Specimens were processed for two-week study, after one dose of *ProLung™-budesonide* was administered via inhalation.

### Data Analysis

One-way ANOVA with Tukey-Kramer multiple comparison data analysis was used for Mch responses using SigmaStat Statistical Software (Systat software Inc., San Jose, CA). EPO activity analysis was performed using the Student *t* test. Over the Study period, there were no significant variability in inflammation within each group with weekly measurements for all of the inflammatory parameters being evaluated. Therefore, all the weekly measurements are presented as Cumulative data and are presented as mean +/− standard error (SEM). A p<0.05 was considered to be statistically significant for all of the above statistical comparisons.

## ACKNOWLEDGEMENTS

We acknowledge the important contributions of the late Dr. Sandhya Nandedkar, a dedicated researcher, for the development of *ProLung™-budesonide*.

VGSK Technologies, Inc. has utilized the non-clinical and pre-clinical services program offered by the National Institute of Allergy and Infectious Diseases.

This project was funded in part with Federal funds from the Division of Microbiology and Infectious Diseases, National Institute of Allergy and Infectious Diseases, National Institutes of Health, Department of Health and Human Services, under Contract No. N01-AI-30048.

Research was also partially funded by Children’s Foundation Grant, Children’s Hospital of Wisconsin, Milwaukee WI.

## REFERENCES

1. He X, Yao F, Chen J, Wang Y, Fang X, Lin X, Long H, Wang Q, Wu Q. The poor prognosis and influencing factors of high D-dimer levels for COVID-19 patients. Sci Rep. 2021 Jan 19;11(1):1830. doi: 10.1038/s41598-021-81300-w. PMID: 33469072; PMCID: PMC7815913.

2. Grifoni E, Valoriani A, Cei F, Lamanna R, Gelli AMG, Ciambotti B, Vannucchi V, Moroni F, Pelagatti L, Tarquini R, Landini G, Vanni S, Masotti L. Interleukin-6 as prognosticator in patients with COVID-19. J Infect. 2020 Sep;81(3):452–482. doi: 10.1016/j.jinf.2020.06.008. Epub 2020 Jun 8. PMID: 32526326; PMCID: PMC7278637

3. Monteil V, Kwon H, Prado P, Hagelkrüys A, Wimmer RA, Stahl M, Leopoldi A, Garreta E, Hurtado Del Pozo C, Prosper F, Romero JP, Wirnsberger G, Zhang H, Slutsky AS, Conder R, Montserrat N, Mirazimi A, Penninger JM. Inhibition of SARS-CoV-2 Infections in Engineered Human Tissues Using Clinical-Grade Soluble Human ACE2. Cell. 2020 May 14;181(4):905–913.e7. doi: 10.1016/j.cell.2020.04.004. Epub 2020 Apr 24. PMID: 32333836; PMCID: PMC7181998.

4. Yan J, Horng T. Lipid Metabolism in Regulation of Macrophage Functions. Trends Cell Biol. 2020 Dec;30(12):979–989. doi: 10.1016/j.tcb.2020.09.006. Epub 2020 Oct 6. PMID: 33036870. Zou X, Chen K, Zou J, Han P, Hao J, Han Z. Single-cell RNA-seq data analysis on the receptor ACE2 expression reveals the potential risk of different human organs vulnerable to 2019-nCoV infection. Front Med. 2020 Apr;14(2):185–192. doi: 10.1007/s11684-020-0754-0. Epub 2020 Mar 12. PMID: 32170560; PMCID: PMC7088738.

5. Hosseinzadeh MH, Shamshirian A, Ebrahimzadeh MA. Dexamethasone vs COVID-19: An experimental study in line with the preliminary findings of a large trial. Int J Clin Pract. 2020 Dec 17:e13943. doi: 10.1111/ijcp.13943. Epub ahead of print. PMID: 33332726; PMCID: PMC7883081.

6. Ramakrishnan S, Nicolau DV Jr, Langford B, Mahdi M, Jeffers H, Mwasuku C, Krassowska K, Fox R, Binnian I, Glover V, Bright S, Butler C, Cane JL, Halner A, Matthews PC, Donnelly LE, Simpson JL, Baker JR, Fadai NT, Peterson S, Bengtsson T, Barnes PJ, Russell REK, Bafadhel M. Inhaled budesonide in the treatment of early COVID-19 (STOIC): a phase 2, open-label, randomised controlled trial. Lancet Respir Med. 2021 Apr 9:S2213–2600(21)00160-0. doi: 10.1016/S2213-2600(21)00160-0. Epub ahead of print. PMID: 33844996; PMCID: PMC8040526.

7. Finney LJ, Glanville N, Farne H, Aniscenko J, Fenwick P, Kemp SV, Trujillo-Torralbo MB, Loo SL, Calderazzo MA, Wedzicha JA, Mallia P, Bartlett NW, Johnston SL, Singanayagam A. Inhaled corticosteroids downregulate the SARS-CoV-2 receptor ACE2 in COPD through suppression of type I interferon. J Allergy Clin Immunol. 2021 Feb;147(2):510–519.e5. doi: 10.1016/j.jaci.2020.09.034. Epub 2020 Oct 15. PMID: 33068560; PMCID: PMC7558236.

8. Konduri KS, Nandedkar S, Düzgünes N, Suzara V, Artwohl J, Bunte R, Gangadharam PR. Efficacy of liposomal budesonide in experimental asthma. J Allergy Clin Immunol. 2003 Feb;111(2):321–7. doi: 10.1067/mai.2003.104. PMID: 12589352.

9. Wiedermann FJ, Lederer W, Mayr AJ, Sepp N, Herold M, Schobersberger W. Prospective observational study of antiphospholipid antibodies in acute lung injury and acute respiratory distress syndrome: comparison with catastrophic antiphospholipid syndrome. Lupus. 2003;12(6):462–7. doi: 10.1191/0961203303lu413oa. PMID: 12873048.

10. Schousboe P, Wiese L, Heiring C, Verder H, Poorisrisak P, Verder P, Nielsen HB. Assessment of pulmonary surfactant in COVID-19 patients. Crit Care. 2020 Sep 7;24(1):552. doi: 10.1186/s13054-020-03268-9. PMID: 32894160; PMCID: PMC7475719.

11. Nguyen HA, Rajaram MV, Meyer DA, Schlesinger LS. Pulmonary surfactant protein A and surfactant lipids upregulate IRAK-M, a negative regulator of TLR-mediated inflammation in human macrophages. Am J Physiol Lung Cell Mol Physiol. 2012 Oct 1;303(7):L608–16. doi: 10.1152/ajplung.00067.2012. Epub 2012 Aug 10. PMID: 22886503; PMCID: PMC3469587.

12. Jung YY, Nam Y, Park YS, Lee HS, Hong SA, Kim BK, Park ES, Chung YH, Jeong JH. Protective effect of phosphatidylcholine on lipopolysaccharide-induced acute inflammation in multiple organ injury. Korean J Physiol Pharmacol. 2013 Jun;17(3):209–16. doi: 10.4196/kjpp.2013.17.3.209. Epub 2013 Jun 11. PMID: 23776397; PMCID: PMC3682081.

13. Reed, L.J., and H. Muench.“A Simple Method of Estimating Fifty Percent Endpoints. Am J Hyg 27 (1938): 493–98.

14. Gangadharam PR, Ashtekar DR, Flasher DL, Düzgüneş N. Therapy of Mycobacterium avium complex infections in beige mice with streptomycin encapsulated in sterically stabilized liposomes. Antimicrob Agents Chemother. 1995 Mar;39(3):725–30. doi: 10.1128/aac.39.3.725. PMID: 7793880; PMCID: PMC162612.

15. Konduri KS, Nandedkar S, Rickaby DA, Düzgüneş N, Gangadharam PR. The use of sterically stabilized liposomes to treat asthma. Methods Enzymol. 2005;391:413–27. doi: 10.1016/S0076-6879(05)91023-9. PMID: 15721394.

16. Mason RJ, Dobbs LG. Synthesis of phosphatidylcholine and phosphatidylglycerol by alveolar type II cells in primary culture. J Biol Chem. 1980 Jun 10;255(11):5101–7. PMID: 7372627.

17. Wang C, Xie J, Zhao L, Fei X, Zhang H, Tan Y, Nie X, Zhou L, Liu Z, Ren Y, Yuan L, Zhang Y, Zhang J, Liang L, Chen X, Liu X, Wang P, Han X, Weng X, Chen Y, Yu T, Zhang X, Cai J, Chen R, Shi ZL, Bian XW. Alveolar macrophage dysfunction and cytokine storm in the pathogenesis of two severe COVID-19 patients. EBioMedicine. 2020 Jul;57:102833. doi: 10.1016/j.ebiom.2020.102833. Epub 2020 Jun 20. PMID: 32574956; PMCID: PMC7305897.

18. Morris G, Bortolasci CC, Puri BK, Olive L, Marx W, O’Neil A, Athan E, Carvalho AF, Maes M, Walder K, Berk M. The pathophysiology of SARS-CoV-2: A suggested model and therapeutic approach. Life Sci. 2020 Oct 1;258:118166. doi: 10.1016/j.lfs.2020.118166. Epub 2020 Jul 31. PMID: 32739471; PMCID: PMC7392886.

19. Bhattacharya J, Westphalen K. Macrophage-epithelial interactions in pulmonary alveoli. Semin Immunopathol. 2016 Jul;38(4):461–9. doi: 10.1007/s00281-016-0569-x. Epub 2016 May 12. PMID: 27170185; PMCID: PMC5018989.

20. Düzgüneş N, Pretzer E, Simões S, Slepushkin V, Konopka K, Flasher D, de Lima MC. Liposome-mediated delivery of antiviral agents to human immunodeficiency virus-infected cells. Mol Membr Biol. 1999 Jan-Mar;16(1):111–8. doi: 10.1080/096876899294832. PMID: 10332745.

21. Majumdar S, Flasher D, Friend DS, Nassos P, Yajko D, Hadley WK, Düzgüneş N. Efficacies of liposome-encapsulated streptomycin and ciprofloxacin against Mycobacterium avium-M. intracellulare complex infections in human peripheral blood monocyte/macrophages. Antimicrob Agents Chemother. 1992 Dec;36(12):2808–15. doi: 10.1128/aac.36.12.2808. PMID: 1482150; PMCID: PMC245550.

22. Hafner AM, Corthésy B, Merkle HP. Particulate formulations for the delivery of poly(I:C) as vaccine adjuvant. Adv Drug Deliv Rev. 2013 Oct;65(10):1386–99. doi: 10.1016/j.addr.2013.05.013. Epub 2013 Jun 7. PMID: 23751781.

23. Song Y, Tang C, Yin C. Combination antitumor immunotherapy with VEGF and PIGF siRNA via systemic delivery of multi-functionalized nanoparticles to tumor-associated macrophages and breast cancer cells. Biomaterials. 2018 Dec;185:117–132. doi: 10.1016/j.biomaterials.2018.09.017. Epub 2018 Sep 11. PMID: 30241030.

24. Chung YH, Beiss V, Fiering SN, Steinmetz NF. COVID-19 Vaccine Frontrunners and Their Nanotechnology Design. ACS Nano. 2020 Oct 27;14(10):12522–12537. doi: 10.1021/acsnano.0c07197. Epub 2020 Oct 9. PMID: 33034449; PMCID: PMC7553041.

25. La-Beck NM, Gabizon AA. Nanoparticle Interactions with the Immune System: Clinical Implications for Liposome-Based Cancer Chemotherapy. Front Immunol. 2017 Apr 6;8:416. doi: 10.3389/fimmu.2017.00416. PMID: 28428790; PMCID: PMC5382151. George PM, Barratt SL, Condliffe R, Desai SR, Devaraj A, Forrest I, Gibbons MA, Hart N, Jenkins RG, McAuley DF, Patel BV, Thwaite E, Spencer LG. Respiratory follow-up of patients with COVID-19 pneumonia. Thorax. 2020 Nov;75(11):1009–1016. doi: 10.1136/thoraxjnl-2020-215314. Epub 2020 Aug 24. PMID: 32839287; PMCID: PMC7447111.

